# Validation and testing of an in vitro model to study medical treatments for anterior urethral stricture disease: assessing the potential efficacy of phosphodiesterase-4 (PDE4) inhibition and testosterone

**DOI:** 10.64898/2026.05.13.724950

**Authors:** Lola P. Lozano, Michael J. Volk, Carly D. Miller, Jane E. Berg, Chantal Allamargot, Charles H. Schlaepfer, Jane T. Kurtzman, M. Ben Christensen, Jeremy B. Myers, Alexandria M. Hertz, Amanda R. Swanton, Budd A. Tucker, Bradley A. Erickson

## Abstract

**Objective:** To 1) determine the expression and distribution of all PDE4 isozymes (A-D) along the length of the anterior urethra, 2) culture fibroblasts and epithelial cells from healthy and strictured urethras, 3) investigate an *in vitro* model of anterior urethral stricture disease (aUSD), and 4) assess the therapeutic potential of phosphodiesterase-4 (PDE4) inhibitors and testosterone compared to paclitaxel.

**Methods:** The presence and relative abundance of PDE4 isozymes (A-D) was confirmed using immunohistochemistry on 5 male cadaveric urethras. Human urethral fibroblasts (FBs) were cultured from healthy control urethras of patients undergoing vaginoplasty (n=3) and from idiopathic bulbar urethral strictures (L2S1E2) of patients undergoing urethroplasty (n=3). Epithelial cells (ECs) were cultured from a healthy control urethra and two urethral strictures. To investigate a model of aUSD, Control FBs were stimulated with TGFβ1 and compared to Stricture FBs on assays of cell proliferation and expression of genes relevant to aUSD pathophysiology. To test therapeutics, Stricture FBs were treated with the PDE4 inhibitor, roflumilast, testosterone (T), or paclitaxel and compared to Control FBs on the previously mentioned assays and cell viability.

**Results:** PDE4- A, B, and D were detected along the length of the urethra. Expression levels did not differ between urethral regions. TGFβ1 altered proliferation and gene expression in a dose-dependent manner. Roflumilast and T preserved cell viability and proliferation and decreased expression of genes positively associated with auSD.

**Conclusion:** Urethral FBs and ECs can be cultured from healthy and strictured surgical specimens, enabling *in vitro* research. PDE4 inhibitors and T may be non-cytotoxic alternatives or additions to paclitaxel for aUSD.

**Highlights:**

- PDE4 isozymes A, B, and D are expressed in adult anterior urethras
- PDE4 is expressed equally from proximal bulbar to meatal urethra
- Epithelial cells and fibroblasts can be cultured from healthy and stricture urethra
- TGFβ1 may not be an optimal method to model aUSD *in vitro*
- Unlike paclitaxel, roflumilast and testosterone are not toxic to urethral cells

## Introduction

Urethral stricture disease (USD) is a relatively common urologic condition, often presenting with obstructive voiding patterns, and characterized by the narrowing of the urethral lumen secondary to urethra fibrosis from trauma and/or chronic inflammation.^1^ The American Urological Association (AUA) guidelines recommend that short (<2cm) bulbar strictures should be initially managed with an endoscopic procedure such as a direct visual internal urethrotomy or a urethral dilation.^2^ However, the long-term success rates after the endoscopic procedure are between 20-30%.^3^

Historically, it has been suggested that endoscopic failures should be managed with a urethroplasty^2,3^. However, the recent introduction of a paclitaxel-coated urethral dilating balloon (Optilume®) has altered the treatment algorithm slightly as FDA approval for the balloon was specifically for failures^4^. The balloon, initially developed for endovascular stenosis dilation, is thought to work by first inducing urothelial microtraumas that are then inhibited from scarring (and re-stenosing) with the introduction of the paclitaxel found on the balloon’s coating. Paclitaxel is an antiproliferative, antifibrotic agent that ultimately functions as a non-specific, cytotoxic agent. ROBUST III, the proof-of concept, randomized-controlled study, confirmed the Optilume’s® superiority to standard balloon dilation^5^. However, the non-specific nature of the drug, and the concern for the long-term integrity of post-dilation urethral tissue, suggests that dilating balloons coated with alterative drugs that more specifically target pathophysiologic processes underlying aUSD may prove superior treatments.

The broad goal of our research is to study new drugs that may potentially offer better outcomes than paclitaxel in aUSD, though we acknowledge that this endeavor is cost and resource-intensive. To facilitate drug discovery, we believe it is essential to create a model that can efficiently analyze candidates for intraurethral use. Thus, in this study we assessed an in vitro model of aUSD and the effect of two potential therapeutic agents that address pathophysiologic processes associated with aUSD.

## Methods

### Tissue Procurement of Cadaveric Urethras

Anterior urethras were isolated and removed from 5 male cadavers lacking urologic pathology within 24 hours post-mortem after consent was provided. Each anterior urethra was divided into five anatomical regions: proximal bulbar, distal bulbar, penoscrotal, penile, and meatal. Specimens were subsequently formalin-fixed and paraffin-embedded. Multiple 5 um sections from each region were mounted onto microscope slides and prepared for immunohistochemistry.

### Immunohistochemistry (IHC) of Cadaveric Urethras

Tissue sections were deparaffinized and rehydrated before undergoing antigen retrieval at 650 watts then washed and permeabilized with 0.1% Triton (10 minutes) before treatment with quenching solution (Vector Labs) for 20 minutes and blocking buffer (Vector Labs) for 2 hours at room temperature. Tissues were incubated with primary antibodies overnight at 4C. Secondary antibodies were incubated on tissues for 30 minutes. Primary and secondary antibody details shown in **Table 1**. Chromogen-DAB (Vector Labs) was applied and followed with hematoxylin counterstain. Sections were then dehydrated and mounted on glass slides and coverslips.

**Table 1.** Antibody Details.

| Assay | Type | Antibody | Catalogue # | Company | Dilution |
| --- | --- | --- | --- | --- | --- |
| ICC | Primary | α-Vimentin | Ab8978 | Abcam | 1:100 |
| | | α-E-Cadherin | 3195 | Cell Signaling Technology | 1:750 |
| | Secondary | Dk α-Ms AF647 | 31571 | Thermo Fisher Scientific | 1:1000 |
| | | Dk α-Rb AF488 | 21206 | | |
| Western Blot, IHC | Primary | α-Androgen Receptor | 5153 | Cell Signaling Technology | 1:500 |
| | | α-PDE4-A&D | PD4-101AP | Fabgennix | 1:1000 |
| | | α-PDE4-B | MA5-25677 | Thermo Fisher Scientific | 1:1000 |
| | | α-PDE4-C | PA5-83938 | | 1:500 |
| | | α-β-actin | 8227 | Abcam | 1:1000 |
| | Secondary | HRP-linked Hs α-Ms | 7076 | Cell Signaling Technology | 1:2000 |
| | | HRP-linked Gt α-Rb | 7074 | | |

### Image Analysis of Cadaveric Urethras

Slides for analysis were imaged on the Ariol slide scanner (Leica Biosystems). Representative images in **Figure 1** were collected on an Olympus BX41 microscope with a digital camera (SPOT Imaging) using a 40x objective lens. Each tissue section was processed in pairs: a negative control with the secondary antibody only and a positive stain with primary and secondary antibodies. This paired design enabled quantitative fold-change analysis by comparing DAB signal intensity between matched negative and positive sections from the same tissue block. The following number of pairs for each antibody target were available for analysis: 23 pairs for PDE4-A&D, 22 pairs for PDE4-B, and 21 pairs for PDE4-C.

**Figure 1.**
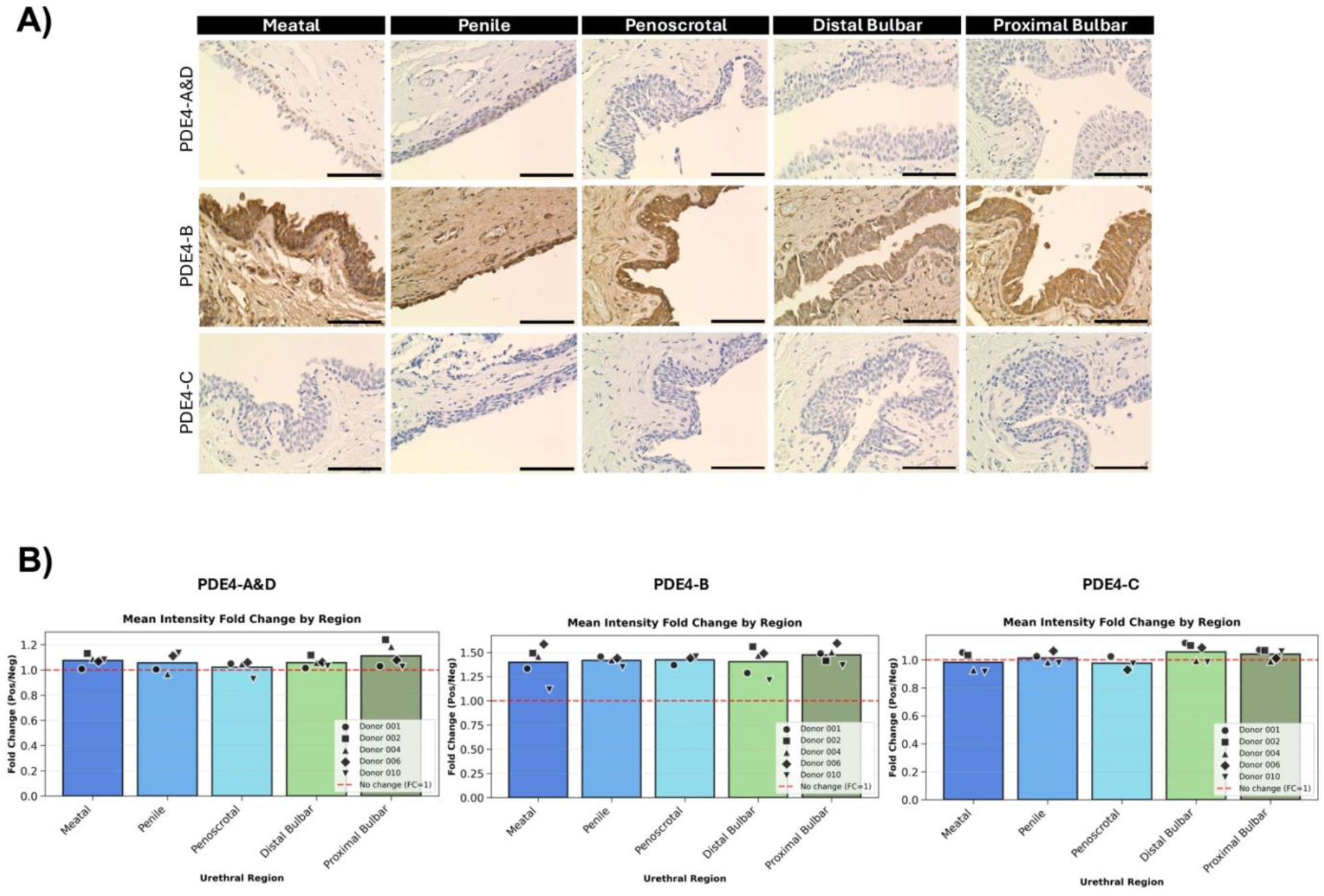
PDE4 isozymes A, B, and D are expressed along the length of the male human anterior urethra. **A)** Representative image from a cadaveric donor lacking urological pathology. Bright field images taken at 40x magnification. Scale bar is 50 um. DAB chromogenic stain for PDE4 isozymes. Hematoxylin counter stain for cell nuclei. **B)** Mean intensity fold change of DAB staining for given antibody targeting PDE4 isozymes A&D, B, or C. Mean intensity between regions for each antibody was compared with ANOVA and found no statistically significant differences.

Briefly, urethral tissue sections were detected on whole slides with converting images to grayscale and applying segmentation and masking. Next, technical artifacts were removed with thresholding to preserve biological signal. Lastly, DAB signal was extracted and quantified. “High DAB” pixels were defined as those exceeding the 75th percentile of DAB intensity within each image, representing the brightest 25% of DAB staining. For each sample, the mean intensity of high DAB pixels was calculated, and fold change was computed as the ratio of positive to negative high DAB mean intensities. Prior to ANOVA, statistical assumptions were verified for each antibody. Normality of residuals was assessed using the Shapiro-Wilk test for each regional group. Homogeneity of variance across regions was evaluated using Levene’s test. Effect sizes were quantified using eta-squared (η^2^), calculated as the ratio of between-group sum of squares to total sum of squares. One-way analysis of variance (ANOVA) was performed separately for each antibody to test for statistically significant differences in mean intensity fold change across the five anatomical regions of the anterior urethra. The significance level (*α*) was set to 0.05 for all statistical tests. Post-hoc pairwise comparisons using Tukey’s Honestly Significant Difference (HSD) test were planned but not performed as no ANOVA results reached statistical significance. All image processing and statistical analyses were performed using Python (version 3.12).

### Tissue Procurement and Culture of Human Urethral Fibroblasts

This study received approval from the University of Iowa Institutional Review Board (IRB No. 202508170). All patients provided informed consent, and research was conducted in accordance with the Declaration of Helsinki. Healthy control urethras (n=3) were isolated from patients undergoing full depth vaginoplasty. Stricture urethras (n=3) were isolated from patients undergoing urethroplasty for idiopathic bulbar urethral strictures > 2cm & ≤ 7cm (L2S1E2).^6,7^ Urethral samples were incubated at 4C overnight in MEMα with Primocin (1:500). Specimens were subsequently washed twice with HBSS with Primocin (1:500) and thoroughly dried then diced with a scalpel into a six-well plate and allowed to dry for an additional 5 minutes before adding media and incubating at 37C, 5% CO2. To culture fibroblasts, these urethral explants received biopsy media (MEMα, heat inactivated fetal bovine serum 1:10, and Primocin 1:500) and were permitted to grow from these urethral explants for ∼2 weeks. To culture epithelial cells, the explants were diced into a six-well plate coated with Matrigel (Corning, Inc.), received Human Urethral Epithelial Cell Media (Cell Applications, Inc.), and were grown until confluent enough to harvest RNA for confirmation of cell identity with qPCR. Media was changed every other day. Fibroblasts were tested for mycoplasma with MycoAlert (Lonza Bioscience) before use in experiments. At passaging, cells were washed with 1xDPBS, lifted with TrypLE (Gibco) for 7 min at 37C, centrifuged in media for 5 min at 200xg, resuspended and counted using Cell Number and Viability on the Moxi GO II (Precision Cell Technologies). Only fibroblasts less than passage 10 were used at a seeding density of 5.0 × 10^4^ cells/ml.

### Immunocytochemistry (ICC) for Confirming Cell Identity of Human Urethral Fibroblasts

Cells were fixed with 4% paraformaldehyde for 20 minutes at room temperature then blocked for 1 hour. Cells were incubated with primary antibody for 2 hours at room temperature or overnight at 4C. Cells were washed with 1xDPBS before incubating with secondary antibody and 1:2000 DAPI (Thermo Fisher Scientific) for 1-2 hours at room temperature. Cells were imaged at 4x and 10x on a Mateo FL Digital Microscope (Leica Microsystems). Primary and secondary antibody details shown in **Table 1**.

### Western Blot for Confirming Expression of Therapeutic Targets in Human Urethral Fibroblasts

Cell pellets were lysed with 75 ul RIPA Lysis and Extraction Buffer with 1:100 Halt™ Phosphatase and Proteinase Cocktail Inhibitors (Thermo Fisher Scientific). Protein concentration was quantified using Pierce™ BCA Protein Assay (Thermo Fisher Scientific) with duplicates of 10 ul per standards and samples. Colorimetric changes in plates were read at 562 nm with the BioTek Cytation 5 microplate reader (Agilent Technologies) after 30 minutes of incubation at 37C. 1.5 ug protein was run on a 4-20% Novex™ Tris-Glycine gel (Thermo Fisher Scientific) for 32 minutes at 226 volts. Broad-range protein transfer to blots was performed using iBlot™ 3 set to a 6 minute run at 25 volts with low cooling (Thermo Fisher Scientific). After incubation with blocking buffer for 1 hour at room temperature or overnight at 4C, blots were probed with primary antibody then washed and probed with HRP-conjugated secondary antibody (as shown in **Table 1**). Blots were imaged using the iBright^™^ FL1500 (Thermo Fisher Scientific) then stripped with Restore^™^ stripping buffer (Thermo Fisher Scientific) and re-probed for additional proteins of interest.

### Pharmacologic Agents

24 hours after plating, cells were stimulated with TGFβ1 or treated with vehicle control, paclitaxel, roflumilast, or testosterone. We chose these latter two drugs given their therapeutic potential modulating subcellular pathways involved in aUSD pathophysiology. Two conditions also characterized by fibrosis and chronic inflammation are chronic obstructive pulmonary disease (COPD) and plaque psoriasis. Notably, while a multitude of drugs are available to treat these conditions, both have FDA-approved drugs that utilize phosphodiesterase-4 (PDE-4) inhibition as means to reverse the fibrotic and inflammatory changes pathognomonic for the condition. These agents act by increasing intracellular cyclic AMP (cAMP), which activates anti-inflammatory and anti-fibrotic pathways by inhibiting TGFβ1 signaling, a key driver of fibrosis in urethral strictures^8^. Importantly, unlike cytotoxic agents, such as paclitaxel, these drugs harness the inherent healing capabilities and may theoretically preserve urethral architecture. Additionally, there is an inverse association between serum testosterone levels and the presence and severity of urethral stricture disease^9-12^. Thus, testosterone as a therapy for aUSD warrants investigation.

### MTS Assay

After 24 hours of stimulation or treatment, 20 ul of CellTiter 96® AQ_queous_ One Solution per well as directed in the manufacturer’s instructions (Promega). After 1.5 hours of incubation with MTS, the absorbance at 490 nm of each well was measured using the BioTek Cytation 5 microplate reader. Change in cell viability was calculated for background subtracted mean values normalized to vehicle control conditions.

### Cell Viability Assay

24 hours post-treatment, percentage of viable cells was determined from single cell suspensions of 250 ul using the Moxi GO II.

### Quantitative Real-Time Polymerase Chain Reaction (RT-qPCR)

cDNA was generated from RNA using SuperScript IV VILO Master Mix (Thermo Fisher Scientific). PCR reactions were run using PrimeTime™ qPCR probes (Integrated DNA Technologies) (**Table 2**) and TaqMan™ Fast Advanced Master Mix (Thermo Fisher Scientific). cDNA was amplified using a QuantStudio 6 Flex Real-Time PCR system (Thermo Fisher Scientific). Data were analyzed using delta-delta Ct method in which values were normalized to the *RPL13A* housekeeping gene and corresponding mRNA values in respective sample control for each assay. Log_2_ fold-change of relative quantification of gene expression was calculated and plotted.

**Table 2.** RT-qPCR Assay Details.

| Relevance to aUSD | Gene | IDT PrimeTime qPCR Assay ID |
| --- | --- | --- |
| Epithelial cell marker | *CDH1* | Hs.PT.58.3324071 |
|  | *KRT19* | Hs.PT.58.4188708 |
|  | *TJP1* | Hs.PT.58.2456962 |
| Mesenchymal cell marker | *FN1* | Hs.PT.58.40005963 |
|  | *ACTA2* | Hs.PT.56a.2542642 |
|  | *COL1A2* | Hs.PT.58.26714610 |
| Increased relative to COL1 in aUSD | *COL3A1* | Hs.PT.58.4249241 |
| Cross links collagen | *LOX* | Hs.PT.58.1471308 |
| Increased in aUSD patients | *CTGF* | Hs.PT.58.4521527.g |
|  | *PDGFB* | Hs.PT.58.45316088 |
|  | *CCL5* | Hs.PT.58.1724551 |
| PDE4 isozyme drug targets | *PDE4A* | Hs.PT.58.40196830 |
|  | *PDE4B* | Hs.PT.58.39087747 |
|  | *PDE4C* | Hs.PT.58.19432716 |
|  | *PDE4D* | Hs.PT.58.2561207 |
| Androgen receptor drug target | *AR* | Hs.PT.56a.14520219 |
| Housekeeping control | *RPL13A* | RPL13A - NM_012423 Set 1 |

### Statistical Analysis

Statistical analysis was performed in GraphPad Prism Version 10.2.3 (347) using one-way ANOVA with Dunnett’s correction (alpha = 0.05) comparing conditions to respective vehicle control condition for each assay. All data were collected from three independent experiments with a case-control pairing.

## Data and Code Availability

The complete analysis pipeline for the cadaveric urethras, including all source code and configuration files, is available at PDE4-Urethra-Analysis. Intermediate analysis outputs are provided for transparency: [artifact detection masks, staining quantification summaries](https://github.com/Mjvolk3/PDE4-Urethra-Analysis/tree/main/notes/assets/images/experiments/002/artifact_analysis), and [individual sample analyses](https://github.com/Mjvolk3/PDE4-Urethra-Analysis/tree/main/notes/assets/images/experiments/002/individual_samples). Raw microscopy images in OME-TIFF format are available upon request as are all other data from cell culture experiments.

## Results

### PDE4 isozymes are expressed along the length of the human male anterior urethra

A single antibody targeted both A and D isozymes of PDE4. It had weak but appreciable staining localized to the epithelium throughout the length of the anterior urethra. PDE4-B stained strongly throughout the length of the urethra, especially within the epithelium, while PDE4-C was undetectable. **Figure 1A** depicts representative images from a single donor along the length of the anterior urethra. One-way ANOVA was performed to assess whether DAB (3,3’-diaminobenzidine) signal intensity varied significantly across the five anatomical regions of the anterior urethra. Two metrics were analyzed for each antibody: mean intensity fold change and high DAB fold change. No significant differences in PDE4 expression were detected between the five anatomical regions (**Figure 1B**).

### Individual cultures of urethral fibroblasts and epithelial cells can be grown from surgical specimens of patients with healthy and strictured urethras

Despite historic difficulty of culturing urethral epithelial cells, we show that it is possible to grow isolated cultures of urethral epithelical cells and fibroblasts from surgical specimens of both healthy and strictured urethras (**Figure 2A**). Allowing cells to grow for ∼4 weeks from explants in HUEpC media at 37C, 5% CO2 enabled selection and proliferation of these cells whose cellular identity was confirmed with RT-qPCR (**Figure 2B-C**). Given that fibroblasts are the cell type responsible for fibrosis and scar tissue formation, we performed our subsequent experiments on these cells after confirming their cellular identity with ICC (**Figure D**).

**Figure 2.**
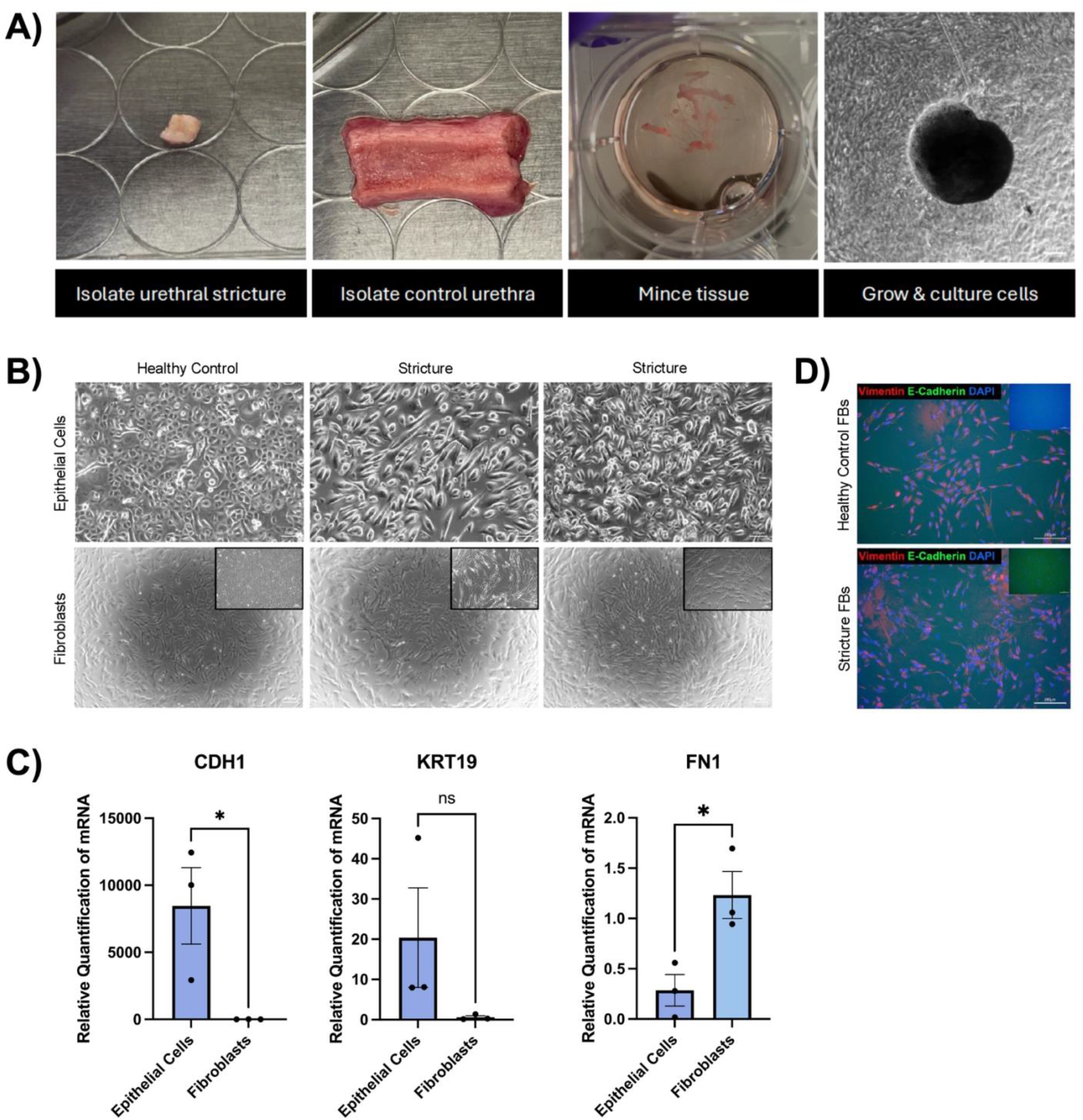
Confirmation of identity of separate cultures of epithelial cells and fibroblasts isolated from the same surgical specimen of healthy and strictured urethras. **A)** Surgical specimens of healthy and strictured urethras can be minced in cell culture dishes for cell growth and culture from tissue explants. **B)** Phase contrast imaging at 4x and 10x showing expected morphology for respective cell type in culture. **C)** RT-qPCR data shows expected increase in epithelial cells markers *CDH1* and *KRT19* compared to fibroblasts and expected increase in *FN1* in fibroblasts relative to epithelial cells. **D)** Immunocytochemistry of cultured urethral fibroblasts confirm expression of cell-specific vimentin and absence of epithelial cell marker E-cadherin. Inset images are negative controls lacking primary antibodies.

### TGFβ1 alters human urethral fibroblast proliferation and gene expression in a dose-dependent manner

Given TGFβ1 as a master regulator of fibrosis and its common application in *in vitro* studies to model fibrosis, we stimulated healthy urethral FBs with a low (2 ng/ml) and high (100 ng/ml) dose of TGFβ1 compared to strictured urethral FBs. While our results showed a dose-dependent trend of decreasing cell proliferation (**Figure 3A**) and increasing expression of genes of interest, none of these results reached statistical significance except for a significant increase in *PDE4C* in healthy cells stimulated with 100 ng/ml TGFβ1 compared to healthy cells treated with vehicle control (p-value = 0.03) (**Figure 3B**).

**Figure 3.**
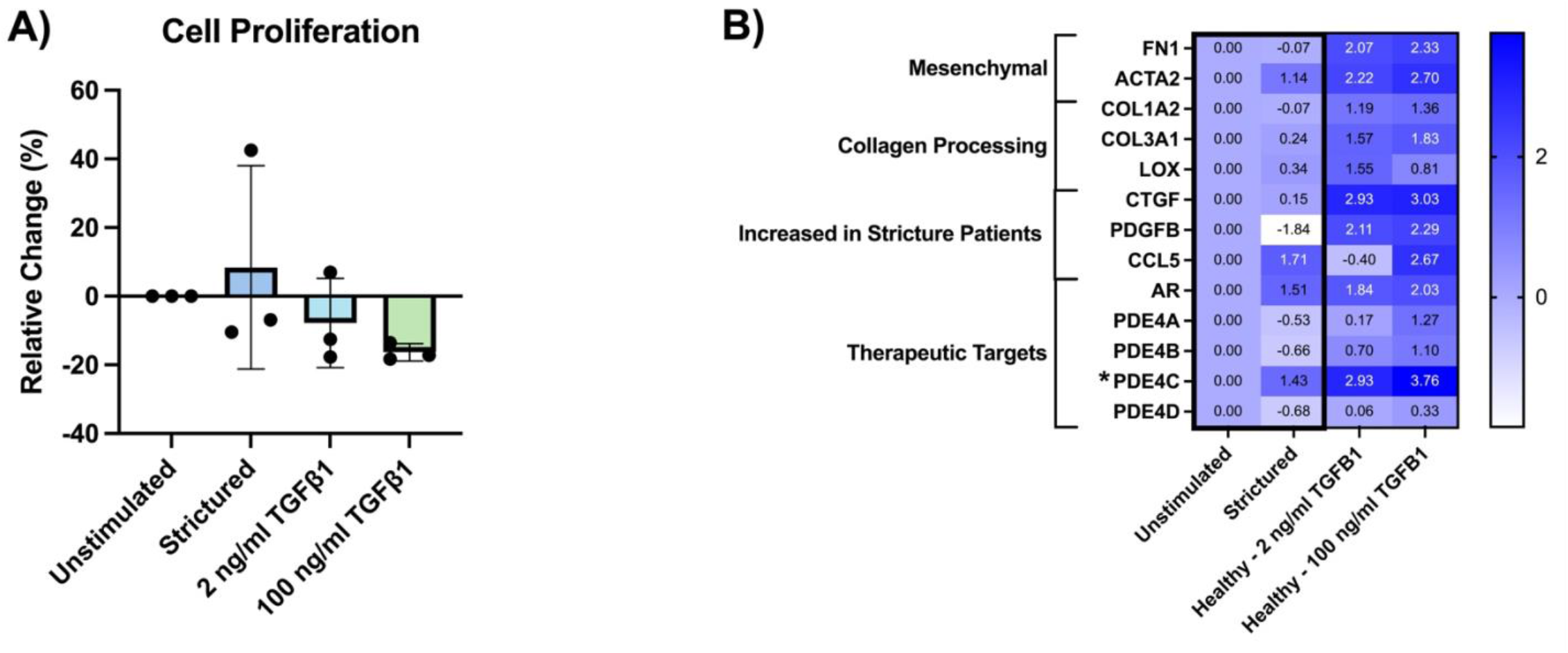
Healthy urethral fibroblasts stimulated with TGFβ1 show a dose-dependent effect on measures of cell proliferation and gene expression. **A)** Cell proliferation decreases with increasing TGFβ1 dosage but does not reach statistical significance. **B)** Log_2_ expression of genes relevant to aUSD pathophysiology increase with increasing TGFβ1 dosage. *; p-value < 0.05.

### Roflumilast and testosterone preserve urethral fibroblast viability and proliferation

We used fibroblasts cultured from urethral strictures to test the therapeutic potential of low and high doses of the PDE4 inhibitor, roflumilast, and testosterone compared to paclitaxel. We first confirmed expression of the targets of these therapeutics at the protein level (**Figure 4A**). While paclitaxel significantly decreased cell viability and proliferation, roflumilast and testosterone left these measures intact (**Figure 4B-C**). There were several significant differences in gene expression when comparing treated stricture cells to healthy cells (**Figure 4D**). High dose roflumilast decreased *PDGF* expression (p-value = 0.03) and high and low doses of testosterone decreased *PDE4D* (p-value = 0.04 and 0.02, respectively).

**Figure 4.**
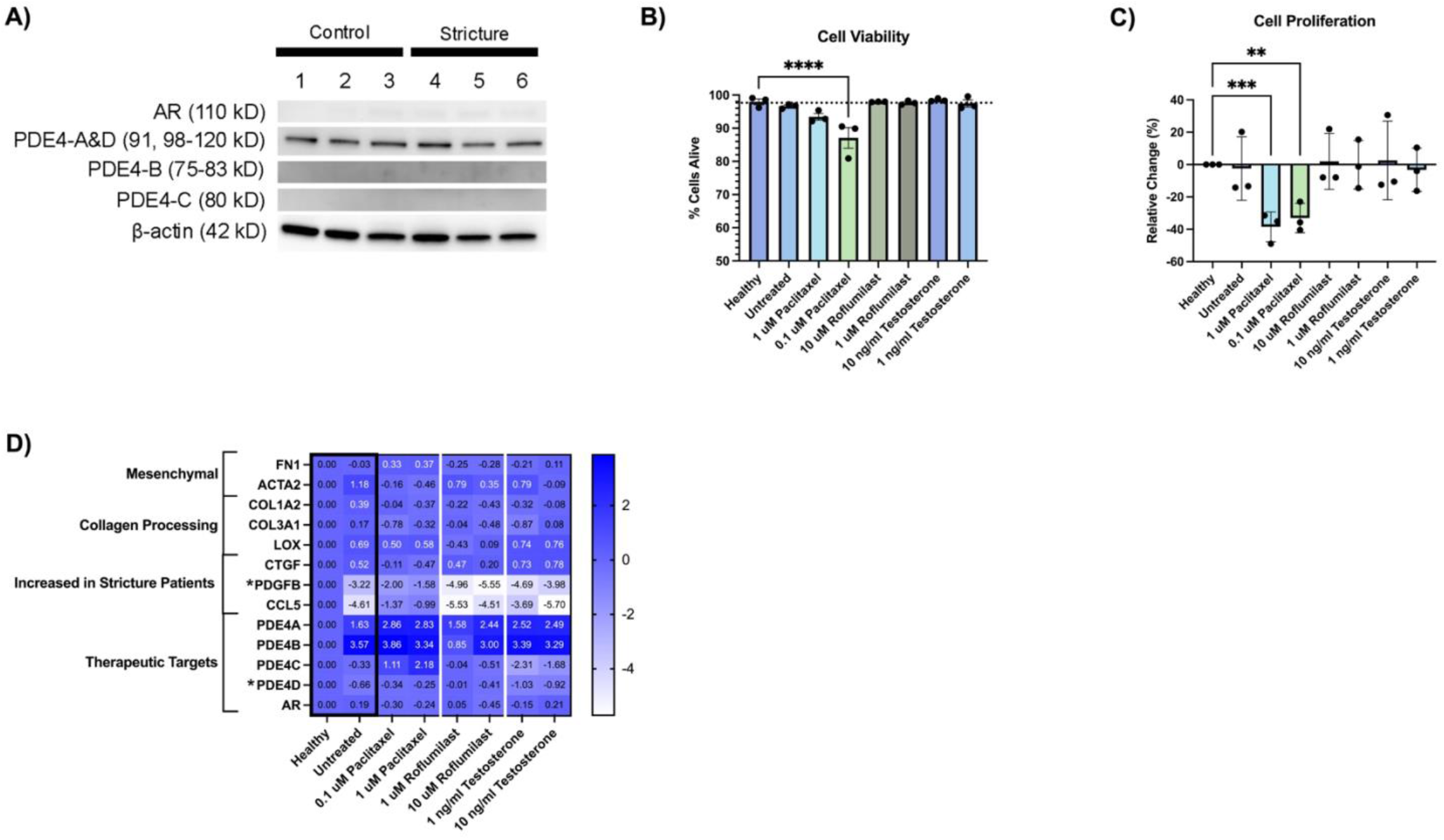
Cultured human fibroblasts from healthy and strictured urethras express therapeutic targets whose effectors preserve cell viability and proliferation. **A)** Western blot for therapeutic targets shows subtle expression of AR and strong expression of PDE4-A&D and house-keeping control β-actin. **B-C)** Paclitaxel decreases cell viability and proliferation. **D)** Aside from *PDGFB* and *PDE4D*, there are no differences in expression of genes involved in aUSD pathophysiology when comparing fibroblasts from healthy urethras versus those from strictured urethras treated with or without low and high levels of drugs.

## Discussion

Due to the high recurrence rate and non-specific cytotoxic mechanism of paclitaxel drug-coated balloons as a treatment for aUSD, we aimed to investigate the therapeutic potential of alternatives in an *in vitro* model of aUSD. We achieved this by first demonstrating that both urethral epithelial cells and fibroblasts can be cultures from explants of surgical specimens of both healthy and strictured urethras. This in turn enables case-control in vitro disease modeling, as well as the ability to test therapeutics on patient-specific cells which can drive a personalized approach to identifying efficient therapeutics for individual patients. Furthermore, healthy urethral fibroblasts can be treated with TGFβ1 to increase expression of genes involved in aUSD pathophysiology.

PDE4 inhibitors and testosterone target many of the processes underlying aUSD pathophysiology (**Figure 5**). Thus, we tested the effect of these drugs on cell viability, proliferation, and gene expression compared to paclitaxel on fibroblasts cultured from idiopathic urethral strictures.

**Figure 5.**
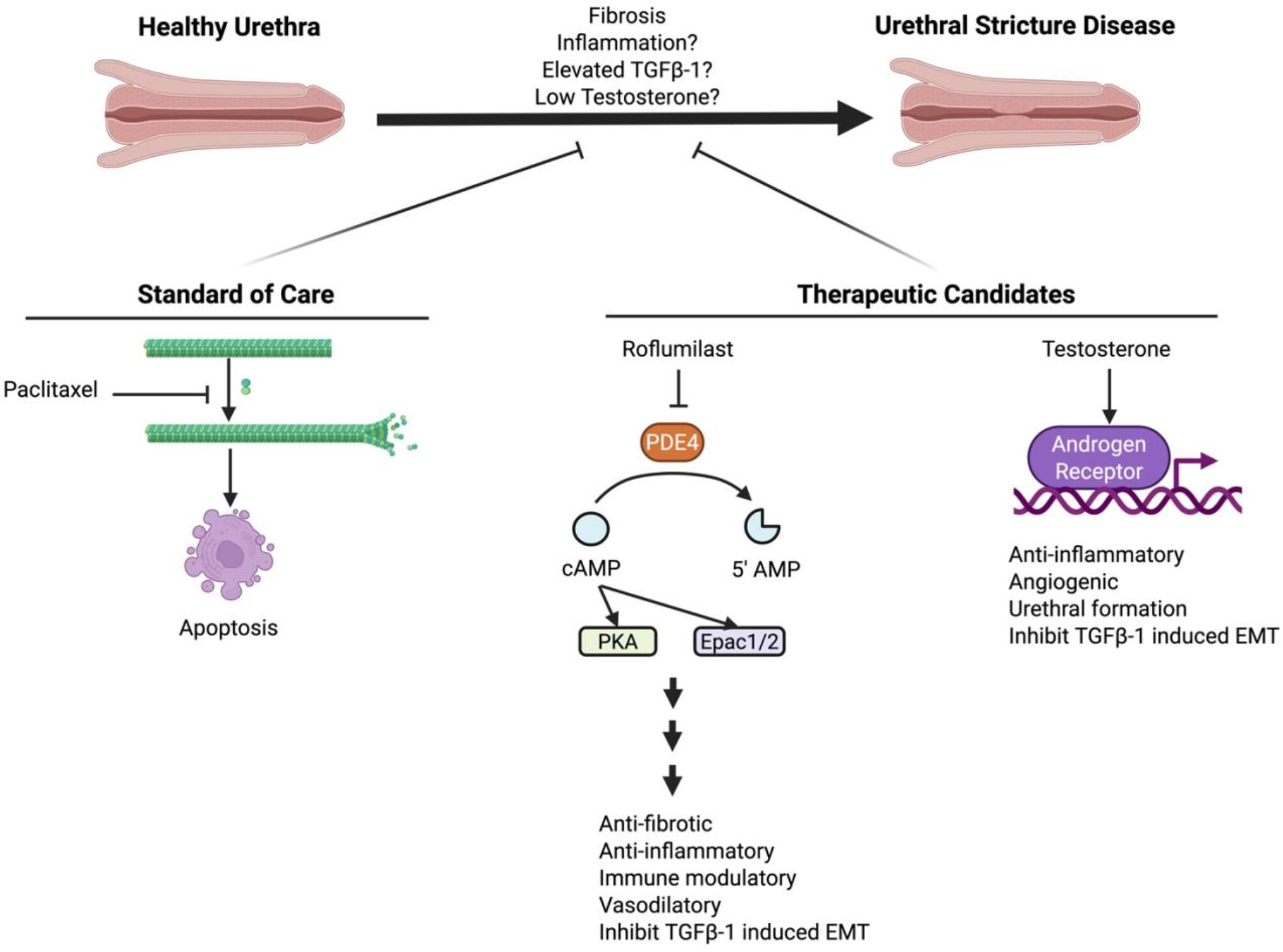
Diagram comparing mechanism of standard of care and alternative therapeutics to target urethral stricture disease. While paclitaxel is the current standard of care delivered on drug-coated balloons to treat urethral stricture disease, it does so by a non-specific mechanism whereby it induces mitotic arrest and causes cell death via apoptosis. Therapeutic candidates like the PDE4 inhibitor, roflumilast, and testosterone may specifically target the pathophysiological processes involved in aUSD.

Identification of homogenous expression of PDE4 isozymes along the anterior urethra was the first step to determining the therapeutic potential of PDE4 inhibitors in aUSD. Prior to this study, the expression of all 4 PDE4 isozymes along the length of the human male anterior urethra was incompletely documented.^1,13-15^ Importantly, this study analyzed the comparative distribution of these isozymes along the length of the human male urethra and included the bulbar urethra which is where the majority of strictures occur.^1^

After confirming homogenous expression of PDE4 isozymes along anterior urethra, we isolated cultures of epithelial cells and fibroblasts from surgical specimens of healthy control urethras and those with idiopathic aUSD. While these cell types have previously been cultured from healthy human urethral surgical specimens, there is no evidence documenting successful culture of epithelial cells from surgical specimens of urethral stricture^16-18^. Thus, researchers must resort to purchasing expensive commercially available cells that are often unavailable for months. We offer urologic translational researchers simplified methods and materials to isolate and culture human urethral epithelial cells from their own surgical specimens, enabling future *in vitro* research on urethral conditions.

Finally, because fibroblasts are the cells primarily involved in aUSD pathogenesis, we compared healthy versus strictured urethral fibroblasts. Cell culture studies often administer TGFβ1 to create a model of fibrosis in a cell type of interest^19^. Studies have produced mixed results regarding the effect of TGFβ1 stimulation on urethral fibroblasts on assays such as cell viability, migration, proliferation, and expression of genes included in our study (e.g. collagens, smooth muscle actin, etc.)^20-23^. While our results did not meet statistical significance, we observed a dose-dependent effect of TGFβ1 on urethral FB proliferation or gene expression. We then used fibroblasts from idiopathic urethral strictures for our experiments assessing the effect low and high dose paclitaxel, the PDE4 inhibitor roflumilast, and testosterone. We found that these drugs decreased expression of genes whose protein products are increased in urethral stricture disease^24-26^. Additionally, we found that unlike paclitaxel, roflumilast and testosterone preserved cell viability and proliferation.

There are several limitations to this study. First, because the same antibody was used to detect PDE4-A&D, we are unable to discern if one or both enzymes are present in our samples. However, given that FDA-approved PDE4 inhibitors are non-specific, we do not anticipate that this difference would affect therapeutic potential. Second, we did not find differences in gene expression between fibroblasts from healthy control and strictured urethras with or without drug treatment. This does not exclude the likelihood of potential differences between these samples, but we may not have detected them because we analyzed a subset of genes hypothesized to play a role in aUSD pathophysiology. Future work using unbiased RNA-sequencing methods on unprocessed surgical samples rather than cultured cells may reveal these differences.

In conclusion, we demonstrated that epithelial cells and fibroblasts can be isolated from surgical specimens of healthy and strictured urethras to model aUSD *in vitro*. While we tested two therapeutic candidates, future work can use this model to investigate the potential of countless single and combination therapies. Our results suggest that compared to paclitaxel, roflumilast and testosterone preserve cell viability and proliferation while reducing expression of genes positively associated with aUSD. Future work is needed to evaluate how PDE4 inhibitors and testosterone might play a role in improving the quality of life of patients living with urethral strictures by improving efficacy and duration of endoscopic procedures and prevent patients from undergoing extensive surgeries and repeat interventions.

## Acknowledgements

We would like to thank the tissue donors and their families for supporting our research. We would like to thank the University of Iowa’s Central Microscopy Research Facility and Dr. Robert Mullins for allowing us access to the microscope in the Chorioretinal Degenerations laboratory.

## Declaration of generative AI and AI-assisted technologies in the manuscript preparation process

During the preparations of this work the authors used ChatGPT in order to assist in manuscript drafting. After using this tool, the authors reviewed and edited the content as needed and take full responsibility for the content of the published article.

